# The role of diets in maternal vaccinations against typhoid toxin

**DOI:** 10.1101/2022.06.10.495645

**Authors:** Durga P. Neupane, Changhwan Ahn, Yi-An Yang, Gi Young Lee, Jeongmin Song

**Affiliations:** Department of Microbiology and Immunology, Cornell University, Ithaca, New York 14853

## Abstract

Children are particularly susceptible to typhoid fever caused by the bacterial pathogen *Salmonella* Typhi. Typhoid fever is prevalent in developing countries where diets can be less well-balanced. Here, using a murine model, we investigated the role of the macronutrient composition of the diet in maternal vaccination efficacies of two subunit vaccines targeting typhoid toxin: ToxoidVac and PltBVac. We found that maternal vaccinations protected all offspring against a lethal-dose typhoid toxin challenge in a balanced, normal diet (ND) condition, but the declined protection in a malnourished diet (MD) condition was observed in the PltBVac group. Despite the comparable antibody titers in both MD and ND mothers, MD offspring had a significantly lower level of typhoid toxin neutralizing antibodies than their ND counterparts. We observed a lower expression of the neonatal Fc receptor on the yolk sac of MD mothers than in ND mothers, agreeing with the observed lower antibody titers in MD offspring. Protein supplementation to MD diets, but not fat supplementation, increased FcRn expression and protected all MD offspring from the toxin challenge. Similarly, providing additional typhoid toxin-neutralizing antibodies to MD offspring was sufficient to protect all MD offspring from the toxin challenge. These results emphasize the significance of balanced/normal diets for a more effective maternal vaccination transfer to their offspring.

**Author summary:** Typhoid fever is a life-threatening systemic infectious disease caused by *Salmonella* Typhi, which is prevalent in developing countries where diets can be less well-balanced. Here, we used mice to study the role of nutrition in maternal vaccination efficacies of two subunit vaccines targeting *Salmonella*’s typhoid toxin. We found maternal vaccinations protected all offspring from a lethal-dose typhoid toxin challenge in a balanced/normal diet (ND) condition, but the lack of protection in a malnourished diet (MD) condition was observed in the PltBVac group. Our data indicate that the difference in maternal vaccination outcomes between ND and MD offspring was due to the less effective maternal antibody transfer from MD mothers to their offspring. Providing additional proteins to MD mothers or additional toxin-neutralizing antibodies to MD offspring saved all malnourished offspring from a lethal-dose typhoid toxin challenge, highlighting the importance of balanced/normal diets for effective maternal vaccination outcomes.

## Introduction

*Salmonella enterica* serovar Typhi (*S*. Typhi) is the cause of the life-threatening systemic disease typhoid fever, resulting in ∼10.9 million reported cases annually and affecting billions of individuals estimated to be exposed to this pathogen [1]. Multi-drug resistant (MDR) and extensively drug-resistant (XDR) *S*. Typhi have emerged, which are spreading globally. Antibiotic-resistant *S*. Typhi is more frequently observed among pediatric patients; more than 90% of the XDR typhoid cases are currently from children younger than 15 years old [2-5]. The typhoid mortality in the pre-antibiotic era was approximately 25% of infected individuals [6]. Two types of typhoid fever vaccines, the live-attenuated Ty21a and Vi subunit vaccines, are currently available [7], whose efficacies are 55∼85%, with the Vi-protein conjugate subunit vaccine being the most efficacious [7-9]. However, there are no vaccines for early-life populations younger than 6 months available [8].

*S*. Typhi is a human-specific pathogen coordinating the expressions of many different virulence factors to establish its infection in humans [10]. Typhoid toxin is a tripartite A_2_B_5_ exotoxin secreted by *S*. Typhi during infection. A pyramid-shaped typhoid toxin consists of CdtB causing host cell DNA damage, PltA ADP-ribosylating host protein(s), and PltB homopentamer binding to a specific glycan receptor moiety expressed on target host cells and tissues [11]. As an exotoxin, typhoid toxin can modulate host cells in the infection site and target cells remotely located to benefit *S*. Typhi [12-14]. In mice expressing “human-type” glycan receptors, typhoid toxin binds to immune cells and brain endothelial cells to intoxicate or pass the toxin to neighbor cells (e.g., neurons). Several typhoid toxins neutralizing IgG and alpaca-derived single domain antibodies have been generated and characterized in detail using mice whose ages were corresponding to human adolescents and adults [15-17].

Newborns are highly susceptible to microbial infections largely due to their immature immune system that is not fully capable of eliciting strong immune responses [18, 19]. Moreover, malnutrition is known to have adverse effects on both innate and adaptive immune responses [20-22], which makes affected individuals more susceptible to microbial infections, including *Salmonella* [22-29]. Maternal vaccinations are proven to be efficacious against flu and pertussis, among others [30-32]. However, little is known about the role of nutrition in maternal vaccination efficacies.

Here, we investigated the role of the macronutrient composition of the diets in maternal vaccination outcomes using two subunit vaccines targeting typhoid toxin as examples in a murine model expressing “human-type” glycans in conjunction with a lethal-dose typhoid toxin challenge. Two primary focuses of these investigations are to determine a correlation between malnutrition and maternal vaccination outcomes and, if any, to find a way(s) to fix the reduced maternal vaccination outcomes by malnutrition.

## Results

### Maternal antibodies protected their progenies from a lethal-dose typhoid toxin challenge

To investigate whether maternal vaccination is effective against typhoid toxin, we vaccinated F1 female mice with either inactive A_2_B_5_ typhoid toxoid (ToxoidVac) or PltB pentamer (PltBVac) twice at weeks 5 and 7 without any supplemental adjuvants, as previously established [14]. These F1 female mice were mated, which resulted in F2 progenies (Fig. 1A). F2 mice were lactated by their F1 mothers for the first 3 weeks, weaned, and examined at week 5 for maternal antibody quantities (Fig. 1A and B). After verifying the presence of anti-typhoid toxin maternal antibodies (Fig. 1B), these F2 mice were challenged with a lethal-dose active typhoid toxin to evaluate whether these maternal antibodies could protect F2 mice from typhoid toxin-mediated illness and death (Fig. 1A, C-F). Maternal antibody titers specific to typhoid toxin subunits available in the plasmas of F2 mice were evaluated by conducting standard end-point enzyme-linked immunosorbent assays (ELISAs). The plasma samples were applied to two types of ELISA plates coated with either A_2_B_5_ typhoid toxin or PltB pentamer to determine antibodies targeting any of the three subunits of typhoid toxin and antibodies specific to receptor-binding PltB subunits, respectively (Fig. 1B). F2 progenies of PBS-administered F1 mothers did not have antibodies specific to typhoid toxin or PltB pentamer, whereas F2 mice maternally-vaccinated with either ToxoidVac or PltBVac exhibited markedly increased maternal antibodies recognizing typhoid toxin subunits (Fig. 1B).

**Figure 1.**
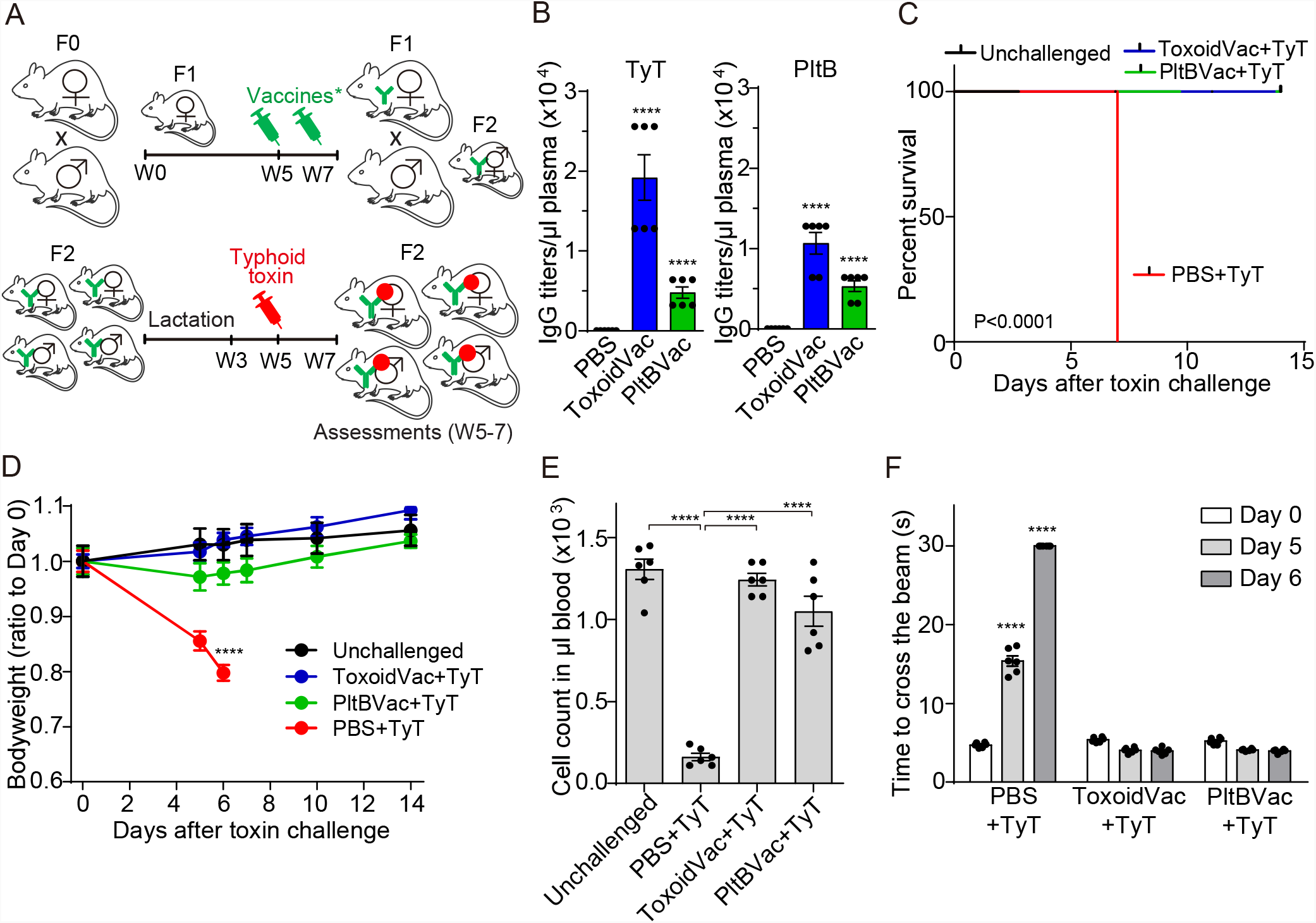
Maternal antibodies protected their progenies from a lethal-dose typhoid toxin challenge. **A**, Schematic cartoon describing the timelines for maternal vaccinations, 2 µg typhoid toxin challenge, and assessments for protection. F1 mothers received two doses (2 µg each) of inactive typhoid toxoid (ToxoidVac) or PltB pentamer (PltBVac) at weeks 5 and 7 (W5 and W7). The vaccinated F1 female mice were mated with unvaccinated F1 male mice, resulting in F2 progenies. Five-week-old F2 mice were challenged with a lethal-dose active typhoid toxin and evaluated for survival, body weight changes, peripheral immune cell counts, and upper motor functions. **B**, Titers of maternal antibodies specific to typhoid toxin (TyT) and PltB pentamer (PltB) available in 1 µl of the indicated F2 mouse plasma collected at week 5. Standard end-point titration methods were used. PBS (black bars), F2 mice from F1 mothers received PBS. ToxoidVac (blue bars), F2 mice maternally vaccinated with ToxoidVac. PltBVac (green bars), F2 mice maternally vaccinated with PltBVac. **C**, Percent survival of the maternally-vaccinated F2 mice challenged with 2 µg typhoid toxin (TyT). **D**, Body weight changes of these F2 mice. **E**, Circulating neutrophil counts in mouse peripheral blood on day 6 after the typhoid toxin challenge. **F**, Balance beam walking results of these F2 mice on days 0, 5, and 6 after typhoid toxin challenge. Bars represent the mean values of two independent experiments ± standard error of the mean (SEM). The total of n=6 per group. ****, p<0.0001, relative to the PBS group for B, to the unchallenged group for D, to the unchallenged group or the TyT group for E as indicated in the graph, and to the day 0 value for F. The log-rank test was performed for C and two-tailed unpaired t-tests for B, D, E, and F.

All of the maternally-vaccinated F2 mice were then challenged with a lethal dose of active typhoid toxin (LD_100_) and analyzed for survival (Fig. 1C), bodyweight changes (Fig. 1D), neutrophil counts (Fig. 1E), and upper motor function defects (Fig. 1F), as previously established [13, 14]. After the toxin challenge, all unvaccinated F2 mice died and showed body weight loss, neutropenia, and upper motor function defect (Fig. 1C-F), as previously demonstrated [13, 14]. In contrast, all maternally-vaccinated F2 mice survived and showed no significant clinical signs for all assessments including body weight changes, leukocyte counts, and neurological signs (Fig. 1C-F), indicating that maternal antibodies transferred to F2 progenies were sufficient to protect them from a lethal dose challenge of active typhoid toxin.

### Maternal antibodies protected some but not all progenies against typhoid toxin

Malnourishment makes affected individuals more susceptible to *Salmonella* infection [21-29]. We next investigated whether maternal vaccinations using ToxoidVac and PltBVac offered protection to malnourished F2 progenies from typhoid toxin-induced illness and death. To establish a malnourished maternal vaccination model, F1 mothers were fed with the control/normal diet (ND) and malnourished isocaloric diet (MD) for a month (weeks 3-7), mated, and continued to feed them with indicated diets throughout the experimental timeframe (Fig. 2A and Table 1). The F2 mice were also fed with indicated diets that matched the diets used for their mothers (Fig. 2A).

**Figure 2.**
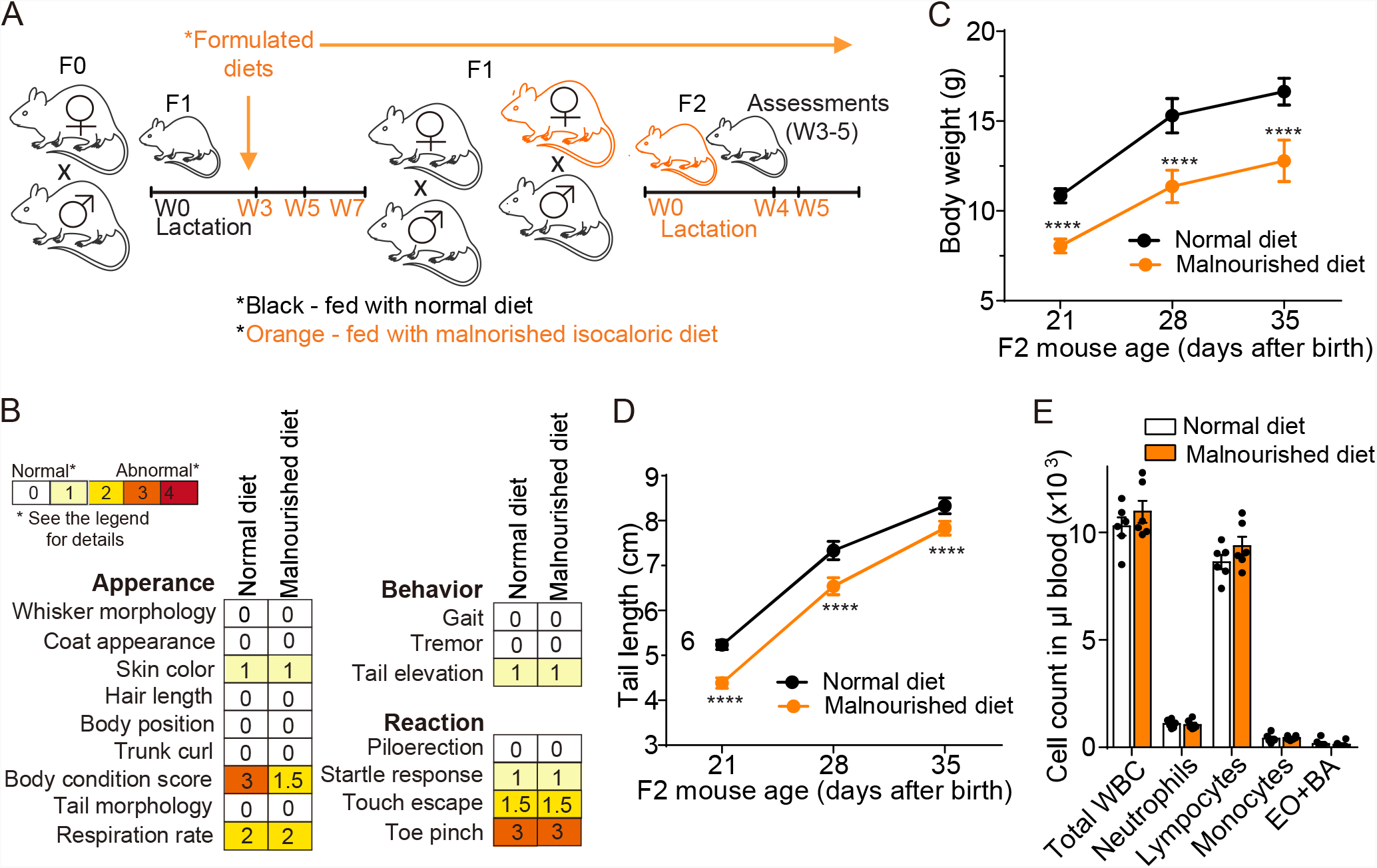
The establishment of a malnourished maternal vaccination model. **A**, Timelines of normal-and malnourished isocaloric diet-feeding and mating that were used in this study. Both F1 and F2 mice were fed normal and malnourished diets. See Table 1 for details. **B**, SHIRPAs on the 5 weeks-old F2 mice. See Table 2 for details. **C**, Bodyweight changes of normal and malnourished diet-fed F2 mice from week 3 to 5. **D**, Tail length changes of these F2 mice. **E**, Circulating leukocyte counts available in 1 µl of peripheral blood of 5 weeks-old F2 mice. WBC, white blood cells. EO, eosinophils. BA, basophils. Lines and bars in C and D represent the mean ± standard deviation (SD), while the mean ± SEM is used for E. n=13-14 F2 mice per group for B-D. n=6 F2 mice per group for E. ****, p<0.0001, relative to normal diet-fed F2 for C and D. Two-tailed unpaired t-tests.

We conducted a standardized mouse phenotype assessment (SHIRPA) on 5-week-old F2 mice and found comparable mouse appearance, behavior, and reaction between ND and MD mice, except for well-conditioned *vs*. skinnier/under-conditioned body appearance (Fig. 2B and Table 2). These results agree with the fact that MD is isocaloric to ND, although it contains less protein and fat (Table 1). Specifically, MD F2 mice exhibited significantly lower body weight and shorter tail length than their ND counterparts (Fig. 2C and D). Notably, immune cell counts were comparable between ND and MD F2 mice (Fig. 2E), indicating that despite a conventional undernourished appearance, MD F2 mice were healthy and had comparable numbers of immune cells to ND F2 mice.

With this animal model, we next investigated whether maternal vaccinations could also protect malnourished progenies from typhoid toxin-induced illness and death. Both ND and MD F2 mice were challenged at week 5 with a lethal-dose typhoid toxin and analyzed for survival, body weight changes, leukocyte counts, and upper motor function defects (Fig. 3A). Similar to Fig 1 where mice were fed with a conventional rodent diet, all ND F2 mice maternally immunized with either ToxoidVac or PltBVac were protected from a lethal-dose typhoid toxin challenge (Fig. 3B-E). However, some but not all MD F2 mice were protected from the typhoid toxin challenge, as 30-40% of MD F2 maternally vaccinated with PltBVac died and showed clinical signs, including statistically significant body weight changes, leukopenia, and upper motor function defects (Fig. 3B-E).

**Figure 3.**
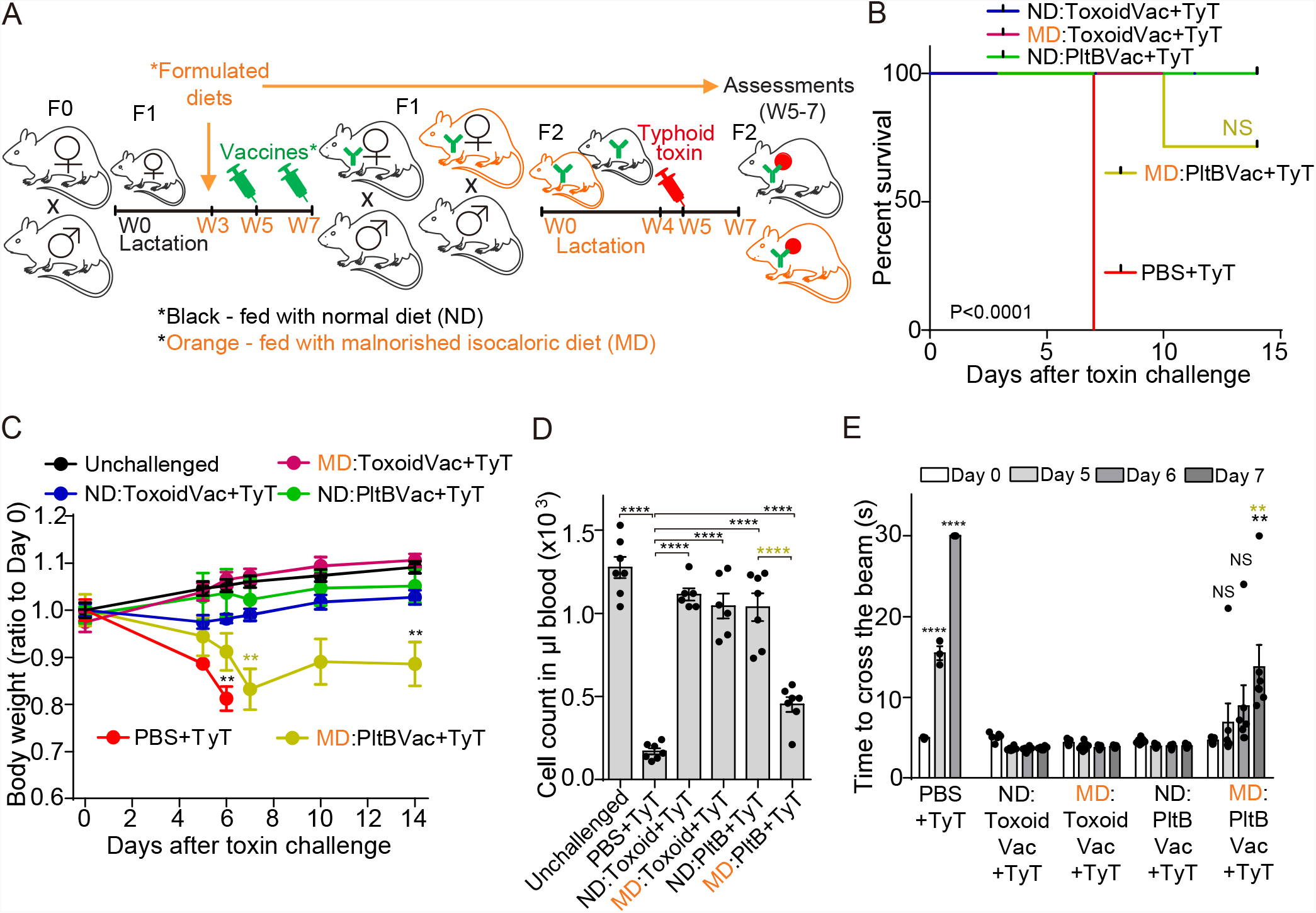
Maternal antibodies protected some but not all progenies against typhoid toxin. **A**, Timelines of indicated diet-feeding, maternal vaccinations, toxin challenge, and assessments for protection. Both F1 and F2 mice were fed with indicated diets, which were started when F1 mice were 3 weeks old and continued throughout the experimental timeframe. **B**, Survival of the maternally-vaccinated F2 mice challenged with 2 µg typhoid toxin (TyT). ND:ToxoidVac (blue), normal diet-fed F2 mice maternally vaccinated with ToxoidVac. MD:ToxoidVac (plum), malnourished diet-fed F2 mice maternally vaccinated with ToxoidVac. ND:PltBVac (green), normal diet-fed F2 mice maternally vaccinated with PltBVac. MD:PltBVac (olive), malnourished diet-fed F2 mice maternally vaccinated with PltBVac. **C**, Body weight changes of these F2 mice after receiving 2 µg typhoid toxin for the 14 day-experimental period. **D**, Circulating neutrophil counts in peripheral blood of these F2 mice on day 6 after the toxin challenge. **E**, Balance beam walking results of these F2 mice on days 0, 5, 6, and 7 after the toxin challenge. Bars represent the mean ± SEM. n=6-7 F2 mice per group. NS, not significant; **, p<0.01; ****, p<0.0001, relative to the unchallenged group for C, to the unchallenged group or the TyT group for D as indicated in the graph, and to the day 0 value for E. Green symbols are used to indicate the statistic analysis results between MD:PltB+TyT group and ND:PltB+TyT group in B-E. The log-rank test was performed for B and two-tailed unpaired t-tests for C, D, and E.

### Unlike F1 mothers, MD progenies had a significantly lower level of typhoid toxin neutralizing antibodies

We hypothesized that malnutrition adversely affects the function and/or titer of typhoid toxin neutralizing antibodies, which resulted in the observed decreased protection in the malnourished condition in the PltBVac group. To examine whether the neutralizing function of antibodies is decreased by malnutrition, we established an *in vitro* infection model using an antibiotic-resistant *S*. Typhi strain to reflect infection of MDR or XDR *S*. Typhi prevalent in endemic countries (Fig. 4A). Human intestinal epithelial Henle-407 cells were infected with *S*. Typhi carrying a kanamycin-resistant plasmid at a multiplicity of infection (m.o.i.) of 30 (rod-shaped red signal in Fig. 4A), where typhoid toxin was continuously secreted by *S*. Typhi(pKan) during culture in the presence of kanamycin (green puncta in Fig. 4A). The secreted typhoid toxin can enter into infected cells and uninfected host cells that can be remotely located [11-14]. After toxin entry into host cells, CdtB of typhoid toxin causes host cell DNA damage that triggers DNA damage repair responses, which can be evaluated by quantifying pH2AX (Ser139) [15] (Fig. 4B). Consistently, wild-type (WT) typhoid toxin triggered a marked increase in the phosphorylation at Ser139 of H2AX, whereas little to no pH2AX signal was observed in cells left uninfected or infected with *S*. Typhi carrying the CdtB^H160Q^ catalytic mutant (Fig. 4B and C), indicating that this *in vitro* infection model can quantitatively evaluate the neutralizing function of antibodies targeting typhoid toxin. To evaluate the neutralizing function of antibodies, 0.01 µl or 0.1 µl plasma samples of indicated F2 mice were added to cell culture media for 24 hours until cells were fixed for microscopy. We found that like all other samples, MD F2 mice maternally vaccinated with PltBVac also neutralized typhoid toxin (Fig. 4B and C).

**Figure 4.**
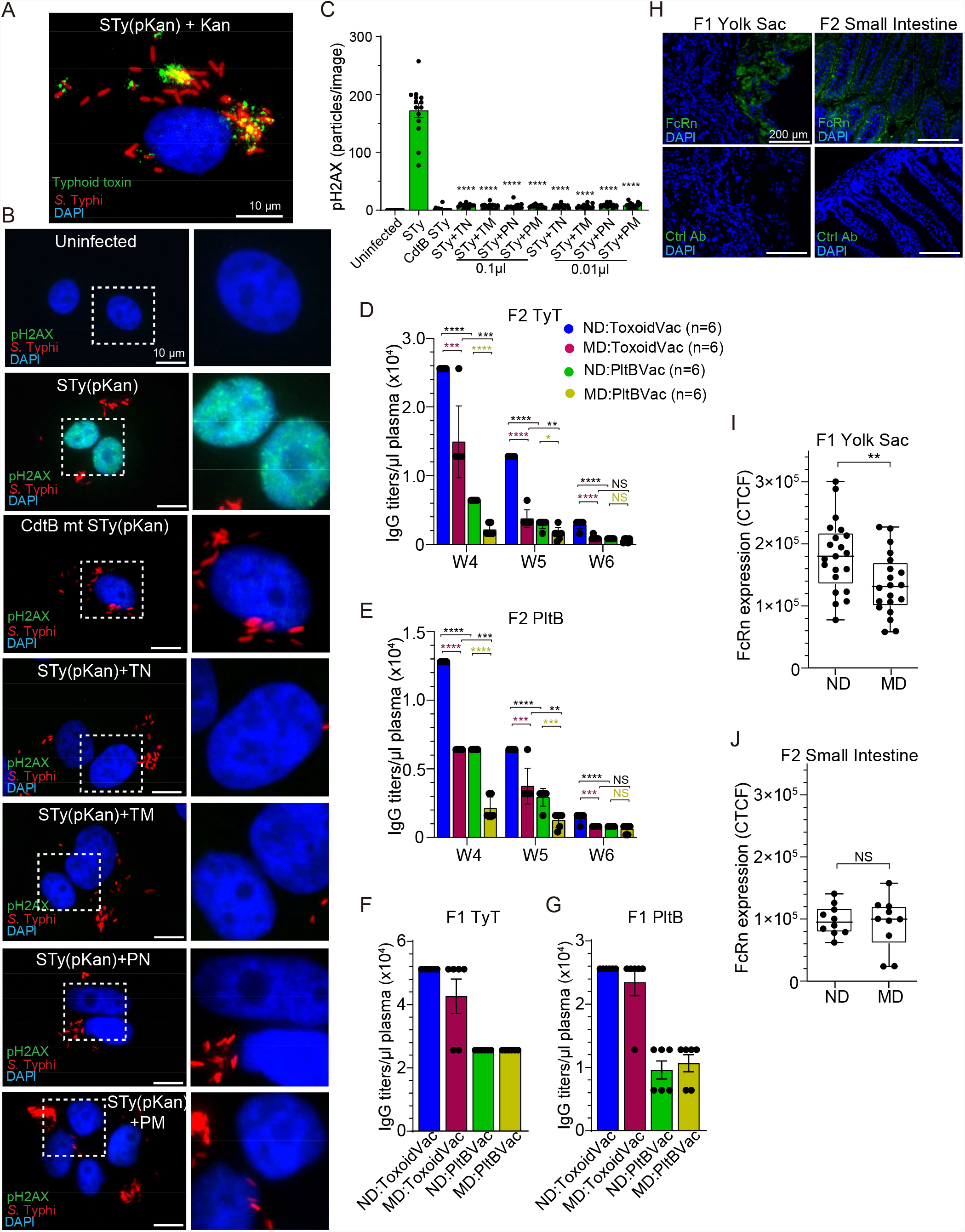
Unlike F1 mothers, MD progenies had a significantly lower level of typhoid toxin neutralizing antibodies. **A**, Representative fluorescence microscope image showing typhoid toxin (green) secreted by *S*. Typhi (red) in kanamycin-resistant *S*. Typhi-infected host cells (blue for host cell DNA). STy(pKan), *S*. Typhi carrying the pKan plasmid. Kan, 50 μg/mL kanamycin treatment. Scale bar, 10 μm. **B**, Representative fluorescence microscope images showing pH2AX (green, reflecting host cell DNA damage repair response), *S*. Typhi(pKan) (red), and host cell DNA (blue). Zoom-in images (right panels) correspond to the dotted boxes in the overall images (left panels). Scale bar, 10 μm. **C**, pH2AX signal quantification of microscopy images obtained from three independent experiments. Bars represent average ± SEM. **** p<0.0001, relative to the STy(pKan) group. Two-tailed unpaired t-tests. **D-E**, Maternal antibody titers specific to typhoid toxin (TyT) (D) and PltB pentamer (PltB) (E) available in 1 µl plasma samples of indicated F2 mice from weeks 4 to 6. ND:ToxoidVac (blue), normal diet-fed F2 mice maternally vaccinated with ToxoidVac. MD:ToxoidVac (plum), malnourished diet-fed F2 mice maternally vaccinated with ToxoidVac. ND:PltBVac (green), normal diet-fed F2 mice maternally vaccinated with PltBVac. MD:PltBVac (olive), malnourished diet-fed F2 mice maternally vaccinated with PltBVac. n=6 per group. **F-G**, IgG titers specific to typhoid toxin (TyT) (F) and PltB pentamer (PltB) (G) that were available in 1 µl plasma samples of the indicated F1 mothers (7 weeks old). n=6 per group. **H**, Verification of the specificity of an anti-FcRn antibody used. Blue, DAPI to stain the nuclei of host cells, Green, FcRn, or isotype control IgG. Scale bar, 200 µm. The placenta and yolk sac tissue were harvested from ND-fed F1 on pregnancy/gestation day (GD)17. The small intestine of F2 was harvested on day 7. **I**, The expression of FcRn on the yolk sac of ND and MD F1 mice. **J**, The expression of FcRn on the small intestine of ND and MD F2 mice. Data from three independent experiments. Boxes and whiskers in I and J represent average, min, and max with all data points. NS; not significant; **, p<0.01. Two-tailed unpaired t-tests.

Next, we evaluated the titers of typhoid toxin neutralizing antibodies available in F1 and F2 mice (Fig. 4D-G). MD F2 mice maternally vaccinated with PltBVac had much lower antibody titers even from week 4, which was further decreased at weeks 5 and 6 (Fig. 4D-E), which is consistent with the lack of protection in some malnourished mice in the PltBVac group (Fig. 3). We found that malnutrition also decreased maternal antibody titers in the ToxoidVac group, supporting the concept that malnourishment-mediated decreased maternal antibodies in F2 progenies may be a universal phenotype for IgGs.

Surprisingly, however, we observed comparable titers of typhoid toxin neutralizing antibodies between ND and MD F1 mothers (Fig. 4F-G), indicating that the reduced maternal antibody titers in MD F2 progenies were largely due to a less effective antibody transfer from MD F1 mothers to their offspring. Because the transfer of maternal antibodies to offspring occurs via the neonatal Fc receptor (FcRn) on the placenta (the yolk sac in the case of mice) during the third trimester of pregnancy and the small intestine during lactation [33, 34], we next evaluated the expression of FcRn on the yolk sac of F1 mothers and small intestine of F2 offspring (Fig. 4H-J). We first verified the specificity of an FcRn antibody (green signal in Fig. 4H); no signal was detected when FcRn was switched with an isotype antibody (bottom panels in Fig. 4H). We probed the placenta tissues attached to the yolk sac of ND and MD F1 mothers and the small intestines of ND and MD F2 offspring with this FcRn antibody. We found that MD mothers expressed a reduced level of FcRn on the yolk sac compared to ND mothers (Fig. 4I), whereas FcRn expression on the small intestines was comparable between ND and MD offspring (Fig. 4J), suggesting that the observed lower antibody titers in MD offspring (Fig. 4D-E) are likely due to a less effective antibody transfer via the placenta/yolk sac during pregnancy.

### MD-induced gut microbiome alterations had no effects on maternal vaccination outcomes against typhoid toxin

Next, we investigated whether this effect is dependent on changes in the gut microbiome that occur as a result of dietary intervention. Through metagenomics of small intestine fecal matters, we found a significant difference in gut microbiome compositions between ND and MD mice, which was further dysregulated by the daily treatment of an antibiotic cocktail consisting of ampicillin, metronidazole, neomycin, and vancomycin (Fig. 5A). We investigated whether this MD-induced gut microbiome alteration contributed significantly to the observed, decreased maternal antibody transfer to their offspring and associated vaccination outcome. We treated ND and MD F1 mice with the above antibiotic cocktail daily via oral gavage for approximately 3 weeks until F1 mothers looked visually pregnant (approximately gestation day 10). This broad-spectrum antibiotic cocktail acts on various aerobic, anaerobic, gram-positive, and gram-negative bacteria; Fig. 5A samples were harvested on gestation day 19. We found that antibiotic treatment did not result in any significant changes in the outcomes of survival (Fig. 5B), bodyweight changes (Fig. 5C), motor function defects (Fig. 5D), typhoid toxin neutralizing antibody titers in F1 mothers (Fig. 5E), and maternal antibody transfer from F1 to F2 progenies (Fig. 5F), indicating that the decreased maternal antibody transfer from F1 to F2 in the malnourished condition is independent of the malnourished diet-mediated gut microbiome.

**Figure 5.**
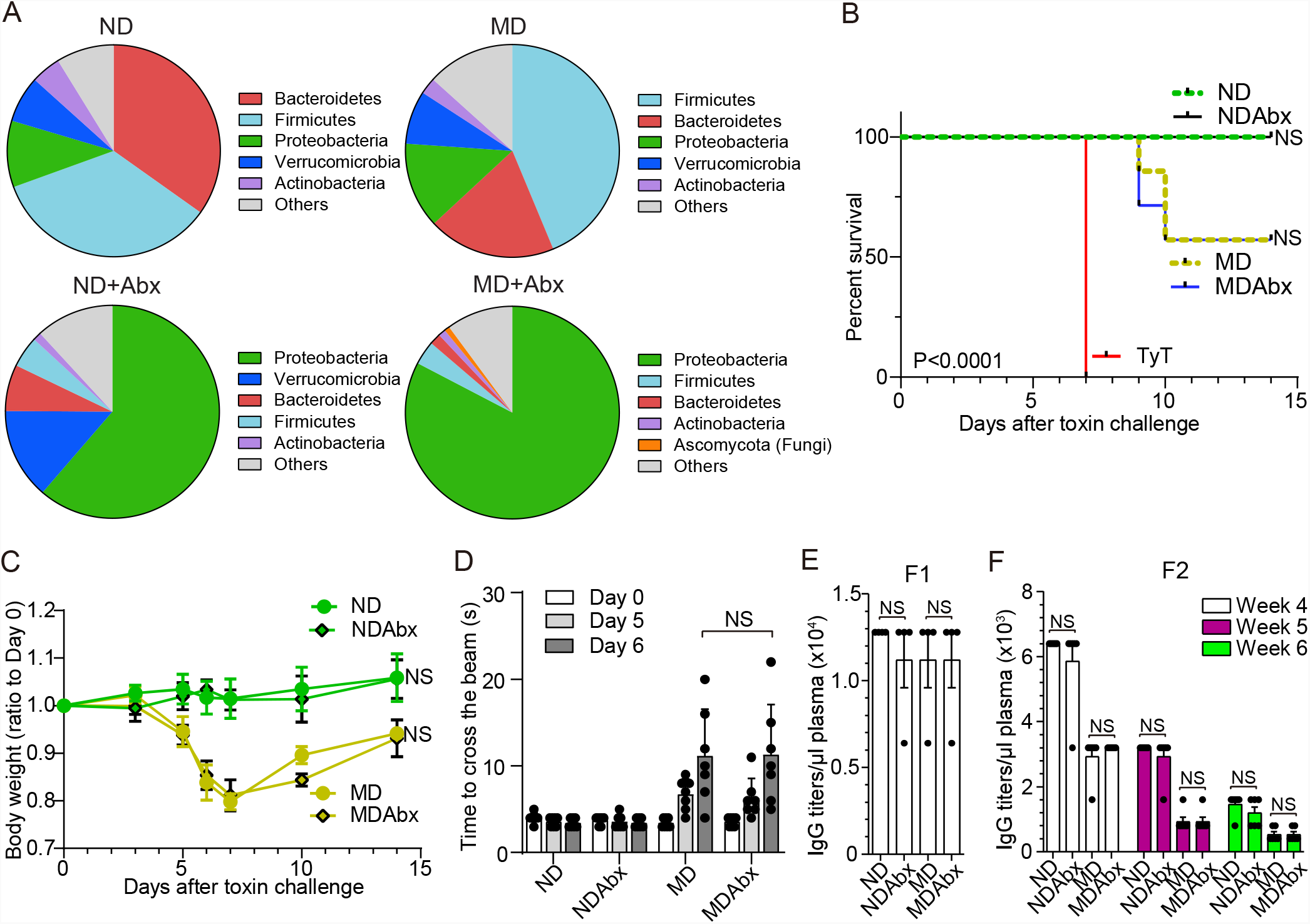
MD-induced gut microbiome alterations had no effects on maternal vaccination outcomes against typhoid toxin. **A**, The small intestine microbiome compositions of ND (n=3), MD (n=3), ND+Abx (n=2), and MD+Abx (n=2). See also Supplementary Table 1. **B**, Survival of the maternally vaccinated (PltBVac) F2 mice challenged with 2 µg typhoid toxin (TyT). ND (green dotted), normal diet-fed F2 mice. NDAbx (black solid), normal diet-fed F2 mice obtained from corresponding antibiotics treated F1. MD (olive dotted), malnourished diet-fed F2 mice. MDAbx (blue solid), malnourished diet-fed F2 mice obtained from corresponding antibiotics treated F1. TyT (red solid), standard regular food-fed F2 mice derived from unvaccinated mothers. **C**. Relative body weight changes of these F2 mice after receiving 2 µg typhoid toxin for the 14 day-experimental period. **D**. Balance beam walking results of these F2 mice on days 0, 5, and 6 after the toxin challenge. **E**. IgG titers specific to PltB pentamer available in 1 μl plasma samples of the indicated F1 mothers (19 days of pregnancy). **F**. Maternal antibody titers specific to PltB pentamer available in 1 μl plasma samples of indicated F2 mice from weeks 4 to 6. The typhoid toxin challenge was conducted in week 5. Lines and bars represent the mean ± SEM. n= 4 (F1) and 6 (F2) mice per group. NS, not significant. The log-rank test was performed for B and two-tailed unpaired t-tests for C-F.

### Providing additional proteins to MD mothers or additional toxin-neutralizing antibodies to MD progenies protected all MD F2 mice from a lethal dose of typhoid toxin challenge

We next tested whether providing additional protein, fat, or both could offer protection to all MD progenies. To supplement protein, fat, or both to MD diets, we added additional 13% calories from protein for protein supplementation, additional 10% calories from fat for fat supplementation, and both for protein and fat supplementation (Table 1, Table 3, and Fig. 6). All other experimental procedures remained the same as before (Fig. 3). Our results indicated that protein supplementation in MD diets, but not fat supplementation, protected all MD progenies from a lethal dose toxin challenge (Fig. 6A-C) and resulted in an efficient maternal antibody transfer (Fig. 6D) and matching FcRn expression on their F1 mothers (Fig. 6E). As expected, similar results were observed when both protein and fat were supplemented to MD, indicating the importance of protein supplementation to MD diets for a more effective maternal antibody transfer from F1 to F2 during pregnancy and subsequent efficacious maternal vaccination outcomes against typhoid toxin.

**Figure 6.**
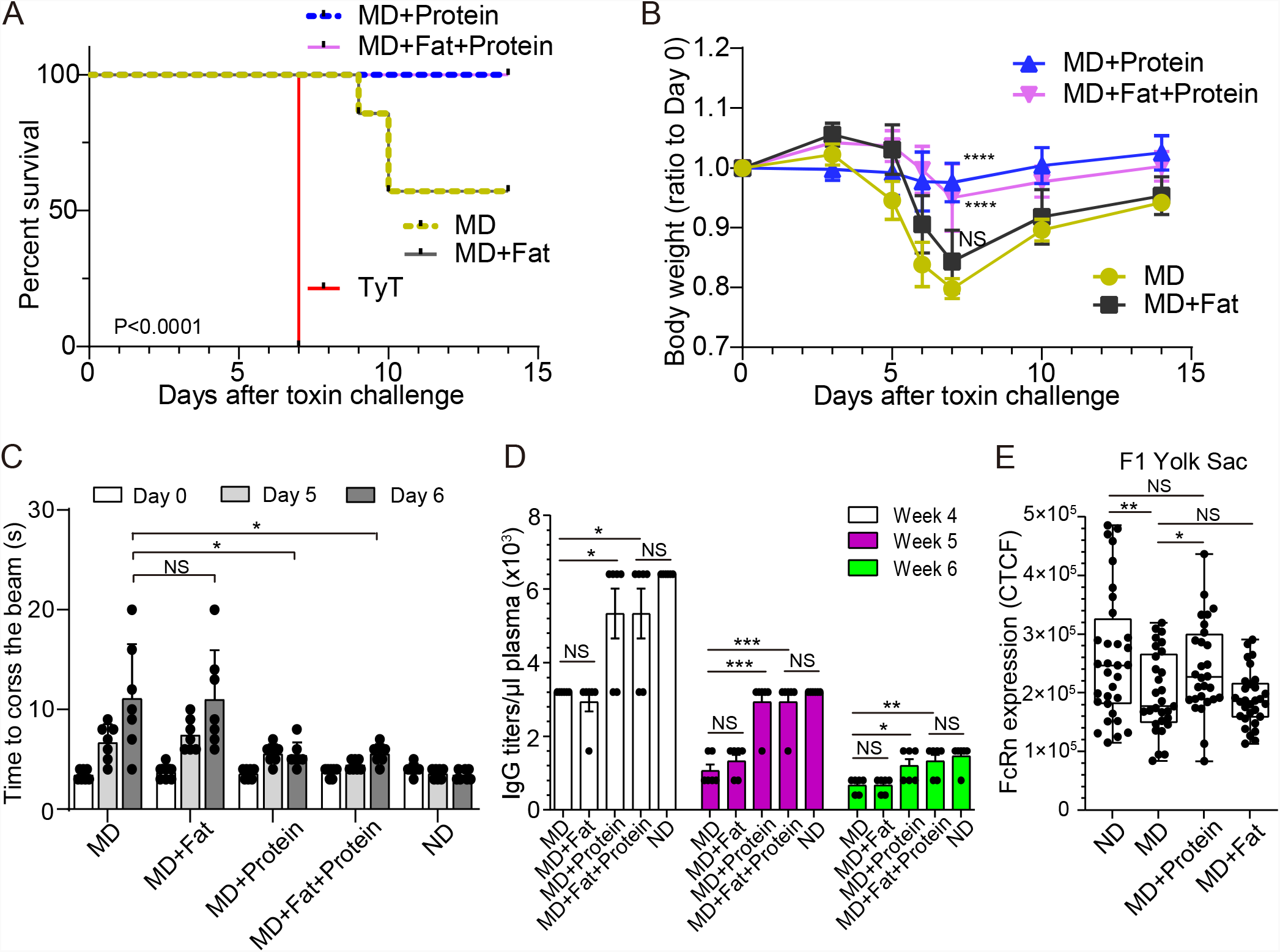
Providing additional protein, but not fat, to MD mothers protected all MD progenies from a lethal dose of typhoid toxin challenge. **A**, Survival of the maternally vaccinated (PltBVac) F2 mice challenged with 2 µg typhoid toxin. All these F2 mice were fed a malnourished diet ± protein/fat supplementation. MD (olive dotted), a malnourished diet alone. MD+Fat (black solid), a malnourished diet supplemented with fat. MD+Protein (blue dotted), a malnourished diet supplemented with protein. MD+Fat+Protein (pink solid), a malnourished diet supplemented with fat and protein. TyT (red solid), standard regular food fed F2 mice derived from unvaccinated mothers. **B**. Relative body weight changes of these F2 mice after receiving 2 µg typhoid toxin for the 14 day-experimental period. **C**. Balance beam walking results of these F2 mice on days 0, 5, and 6 after the toxin challenge. ND (normal diet) fed mice values have been added for reference. **D**. Maternal antibody titers specific to PltB pentamer available in 1 μl plasma samples of indicated F2 mice from weeks 4 to 6. The typhoid toxin challenge was conducted at the beginning of week 5. Lines and bars represent the mean ± SEM. n= 6 F2 mice per group. **E**, The expression of FcRn on the yolk sac of ND, MD, MD+Protein, and MD+Fat F1 mice. Data from three independent experiments. Boxes and whiskers in E represent average, min, and max with all data points. NS; not significant. *, p<0.05; **, p<0.01; ***, p<0.001; ****, p<0.0001 as indicated. The log-rank test was performed for A and two-tailed unpaired t-tests for B-E.

Fig. 6 results led us to further hypothesize that the supplementation of additional neutralizing antibodies directly to MD progenies to ramp up the antibody level above “the threshold” required for protection could protect all MD F2 mice against a lethal dose of typhoid toxin challenge. To test this hypothesis, we supplemented MD F2 mice with anti-PltB IgG TyTx4 or anti-CdtB IgG TyTx11 [15, 17], which resulted in 100% survival, normal body weight changes, neutrophil counts, and motor functions (Fig. 7A-D). These results indicate that malnutrition has an adverse effect on maternal vaccination outcomes against typhoid toxin, which is largely due to a less effective maternal antibody transfer from F1 mothers to their progenies. Therefore, the lack of protection observed in some MD F2 mice can be remedied by providing additional protein to MD mothers to increase the transfer of maternal antibodies to F2 or additional typhoid toxin neutralizing antibodies directly to MD F2 mice when their toxin neutralizing antibody level is below the protection threshold.

**Figure 7.**
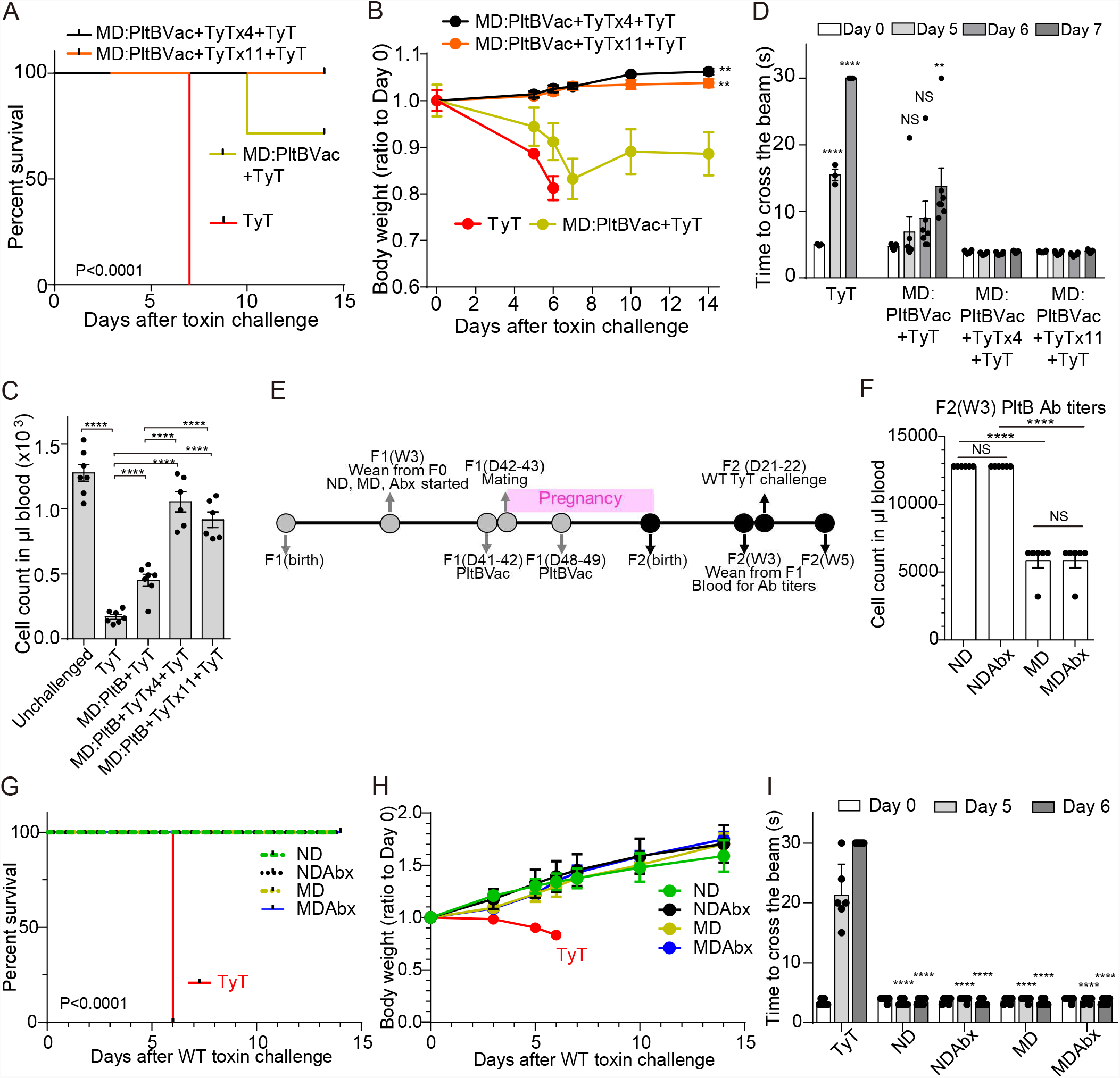
Providing additional toxin-neutralizing antibodies to MD progenies protected all MD F2 mice from a lethal dose of typhoid toxin challenge. **A**, Survival of the F2 mice received 2 µg typhoid toxin (TyT). MD:PltBVac (olive), malnourished diet-fed F2 mice maternally vaccinated with PltBVac. MD:PltBVac+TyTx4 (black), malnourished diet-fed F2 mice maternally vaccinated with PltBVac and administered with TyTx4 (monoclonal antibody recognizing the PltB subunits) 30 min before the typhoid toxin challenge. MD:PltBVac+TyTx11 (orange), malnourished diet-fed F2 mice maternally vaccinated with PltBVac and administered with TyTx11 (monoclonal antibody recognizing the CdtB subunit). n=6-7 F2 mice per group. **B**, Body weight changes of these F2 mice for the 14 day-experimental period after the typhoid toxin challenge. **C**, Circulating neutrophil counts in peripheral blood of these F2 mice on day 6 after the typhoid toxin challenge. **D**, Balance beam walking results of these F2 mice on days 0, 5, 6, and 7 after the typhoid toxin challenge. **E-I**, Maternal antibody titers available in MD-PltBVac F2 mice at W3 were above the threshold antibody level required for full protection. **E**, Timelines of indicated diet-feeding, maternal vaccinations, toxin challenge, and assessments for protection. Both F1 and F2 mice were fed with indicated diets, which were started when F1 mice were 3 weeks old and continued throughout the experimental timeframe. **F**. Maternal antibody titers specific to PltB pentamer available in 1 μl plasma samples of indicated F2 mice at week 3 (W3). The typhoid toxin challenge was conducted 1-day after harvesting blood samples for antibody measurement. **G**, Survival of the maternally vaccinated (PltBVac) F2 mice challenged with 2 µg typhoid toxin (TyT). ND (green dotted), normal diet-fed F2 mice. NDAbx (black dotted), normal diet-fed F2 mice obtained from corresponding antibiotics treated F1. MD (olive dotted), malnourished diet-fed F2 mice. MDAbx (blue solid), malnourished diet-fed F2 mice obtained from corresponding antibiotics treated F1, TyT (red solid), standard regular food-fed F2 mice derived from unvaccinated mothers. **H**. Relative body weight changes of these F2 mice after receiving 2 µg typhoid toxin for the 14 day-experimental period. **I**. Balance beam walking results of these F2 mice on days 0, 5, and 6 after the toxin challenge. Lines and bars represent the mean ± SEM. n= 6 (F2) mice per group. NS, not significant. Bars represent the mean ± SEM. NS, not significant; **, p<0.01; ****, p<0.0001, relative to MD:PltBVac+TyT for B, to the group as indicated in the graph for C and F, to the day 0 values for D, and to the corresponding TyT group for I. The log-rank test was performed for A and G, while two-tailed unpaired t-test was conducted for B-D, F, and I.

Malnourishment resulted in a significant decrease in maternal antibody transfer in both ToxoidVac and PltBVac, although we found that ToxoidVac always induced higher levels of toxin neutralizing antibodies than PltBVac (Fig. 4D-E). These results support the concept that there is a protection threshold level of typhoid toxin neutralizing antibodies. To further investigate this hypothesis, we modified experimental time points to examine maternal antibody titers available in MD F2 mice maternally vaccinated with PltBVac before week 4 (Fig. 7E-I). Consistent with all other findings, an MD-dependent decrease in maternal antibody transfer was observed at week 3, compared to their ND counterparts (Fig. 7F). Also in line with a time-dependent gradual decrease of maternal antibody levels between W4-6 (Fig. 4D-E), we observed at least 2-fold higher antibody titers in 3-week-old F2 mice than their 4-week counterparts in both ND and MD conditions (Fig. 7F *vs*. Fig. 4E). These 2-fold higher antibody titers in MD PltBVac F2 mice at W3 match the titers available in ND PltBVac F2 mice at W4, the group that was protected against a lethal dose of typhoid toxin challenge (Fig. 3B, 4E, 5B-F). In support of the protection threshold antibody level hypothesis, we found that all MD PltBVac F2 mice survived and were protected from typhoid toxin-mediated *in vivo* toxicities when they were challenged at W3 (Fig. 7G-I). Moreover, the antibiotic cocktail was administered daily via oral gavage throughout Fig 7E-I experiments; consistent with Fig 5 results, we found that the effect is independent of the gut microbiome.

## Discussion

Newborns and children are particularly susceptible to typhoid fever. Malnourishment makes this vulnerable population even more susceptible to microbial infections including *Salmonella*. Antibiotic-resistant MDR and XDR *S*. Typhi became dominant in endemic countries, as nearly all clinical isolates of *S*. Typhi are either MDR or XDR. Infection by antibiotic resistance strains often results in more severe clinical manifestations, as they produce their virulence factors even while being treated with antibiotics. As maternal vaccination is one of the most effective intervention strategies for protecting mothers and their offspring [35], we investigated whether maternal vaccinations against typhoid toxin are effective in protecting malnourished progenies from a lethal dose of typhoid toxin challenge.

This study made three important findings. First, we demonstrated that two subunit vaccines, ToxoidVac and PltBVac, without any supplemental adjuvants are efficacious, as they offered protection to all maternally vaccinated newborns against a lethal dose challenge of active typhoid toxin (Fig. 1). The capabilities of these subunit vaccines eliciting strong humoral immune responses without adjuvants are likely associated with the activation of immune cells/antigen-presenting cells by typhoid toxin’s receptor-binding subunit PltB that recognizes a glycan receptor moiety displayed on the surface of antigen-presenting cells [12, 14]. In this regard, our study indicates that the glycan receptor binding PltB of ToxoidVac and PltBVac appears to effectively stimulate humoral immune responses leading to the production of typhoid toxin-neutralizing antibodies. This observation is not too surprising, as some toxoids are being used as adjuvants of human vaccines, such as tetanus toxoid for typhoid vaccine [9], while many others have also been demonstrated to be efficacious against various bacterial, viral, and parasitic pathogens [9, 36-39]. Furthermore, when the same concentrations of these two subunit vaccines were used, ToxoidVac elicited stronger humoral immune responses for anti-PltB antibodies than PltBVac (Fig. 1). The exact mechanism of how additional subunits in ToxoidVac, inactive PltA and CdtB subunits, helped elicit stronger antibody responses to PltB pentamer needs to be investigated in the future, but it is intriguing to speculate that the antigen presentation process may become more productive when the other subunits, particularly inactive CdtB subunit, are present.

Second, we demonstrated that, unlike balanced/normal diet-fed mice, vaccination offered protection only to some malnourished offspring after 1 month after birth (Figs. 2 and 3). We found that this effect is primarily due to a less effective maternal antibody transfer from mothers to their offspring in the malnourishment condition, compared to the ND condition (Figs. 4 and 6E). In particular, PltBVac elicited less strong humoral responses in malnourishment and a reduced transfer of maternal antibodies targeting typhoid toxin (Figs. 3 and 4). We found that malnourished mothers expressed a reduced level of FcRn on the yolk sac compared to balanced/normal diet-fed mothers, whereas FcRn expression on the small intestines was comparable between ND and MD offspring (Fig. 4), suggesting that the observed lower antibody titers in MD offspring are likely due to a less effective antibody transfer via the placenta/yolk sac during pregnancy. Consistently, protein supplementation to MD F1 mothers resulted in a significant increase in FcRn expression, maternal antibody transfer, and protection against typhoid toxin challenge (Fig. 6). Future studies would determine the roles of other factors in the maternal antibody transfer in malnutrition, including antibody isotypes and glycan modifications [40]. Nonetheless, the reduced maternal antibody transfer in malnutrition is in agreement with studies by others [33, 34]. For instance, a low-level placental transfer of maternal antibodies targeting *Haemophilus influenza* type b in malnourished pregnant women was previously observed [41], suggesting that malnutrition-associated adverse effects on maternal vaccinations may be a universal phenomenon against various maternal vaccines. Moreover, using an *in vitro* infection model, we demonstrated that maternal antibodies transferred to malnourished offspring were also functional (Fig. 4), suggesting that malnourished offspring can also be fully protected if the antibody quantities are elevated, which was tested in Figs 6 and 7 experiments.

Third, we demonstrated that providing additional protein to malnourished mothers (Fig. 6) or additional typhoid toxin neutralizing antibodies to malnourished offspring (Fig. 7) is sufficient to save all malnourished F2 mice from a lethal dose typhoid toxin challenge. This is in agreement with the observation that malnutrition negatively affects the transfer of maternal antibodies from pregnant mothers to their progenies.

From a methodology standpoint, we established a murine model to investigate the role of nutrition in maternal vaccination outcomes against typhoid toxin. This model enabled us to distinguish the contribution of protein from fat in maternal vaccination outcomes against typhoid toxin (Fig. 6). This *in vivo* model, along with other *in vivo* and *in vitro* typhoid toxin intoxication models utilized in this study would help quantitatively evaluate other intervention strategies targeting typhoid toxin. Moreover, we predict that the established malnutrition model may serve as a critical resource for other vaccine studies evaluating the role of malnutrition in maternal vaccination outcomes.

In summary, we demonstrated that malnutrition resulted in a less effective maternal antibody transfer from pregnant mothers to their progenies, offering protection to some but not all malnourished offspring. However, providing additional protein to malnourished pregnant mothers or additional typhoid toxin neutralizing antibodies directly to malnourished offspring was sufficient to ramp up the toxin-neutralizing antibody level and protect all malnourished offspring against typhoid toxin. These results emphasize the significance of balanced/normal diets for effective maternal vaccination outcomes.

## Materials and Methods

### Ethics statement

Mouse experiments were carried out in accordance with protocol #2014-0084 approved by Cornell University’s institutional Animal Care and Use Committee, which followed IACUC and AAALAC guidelines.

### Bacterial strains

WT and CdtB catalytic mutant *Salmonella enterica* serovar Typhi ISP2825 have been described previously [11, 42-44]. For infection experiments, *S*. Typhi strains harboring a plasmid carrying a kanamycin-resistant gene were grown at 37 °C in 2 mL LB broth containing 0.3 M NaCl and 50 µg/ml kanamycin to an OD_600nm_ of ∼0.9 after inoculation from an overnight culture in 2 mL LB broth at a dilution of 1:50.

### Mammalian cell culture conditions

Human intestinal epithelial Henle-407 cells were cultured in DMEM high glucose (Invitrogen) and RPMI-1640 (Invitrogen) supplemented with 10% FBS (Hyclone cat# SH30396.03, Lot# AD14962284), respectively. Sialic acid contents of the FBS used were validated, which was ∼99% Neu5Ac and less than 1% Neu5Gc. Cells were kept at 37°C in a cell culture incubator with 5% CO_2_. Mycoplasma testing was conducted regularly as part of the cell maintenance practice.

### Expression and purification of recombinant typhoid toxin, typhoid toxoid, and PltB pentamer

Overexpression and purification of typhoid toxin, typhoid toxoid, and PltB homopentamer were carried out as previously described [13-15, 17, 45].

### *In vivo* mouse experiments

The mice used in this study were originally purchased from the Jackson Laboratory and bred in a vivarium in the ECRF animal facility at Cornell University. Age-and sex-matched C57BL/6 mice were randomly allocated to each group. Experimental schemes along with mouse ages are indicated in each figure. When indicated, mice were fed with either control/normal or malnourished isocaloric diets (Table 1). Mice fed with these formulated diets were evaluated by a modified SHIRPA protocol with an emphasis on appearance, behaviors, and reactions as described previously [14, 46] (Table 2). When indicated, the MD diet was supplemented with protein (additional 13% calories from casein and L-cysteine), fat (additional 10% calories from soybean oil), or both (Table 3). For Fig. 5 experiments, F1 mothers were treated with an antibiotic cocktail consisting of 1 mg/mL ampicillin, 0.5 mg/mL vancomycin, 1 mg/mL neomycin, and 1 mg/mL metronidazole 3 weeks after ND and MD diets were started (for Fig. 5 experiments) or at 3 weeks when ND and MD diets were started (for Fig 7E-I experiments). One hundred microliters of the antibiotic cocktail were administered daily via oral gavage continuously until the mothers look visually pregnant (approximately gestation day 10) for Fig. 5 experiments or throughout Fig 7E-I experiments. All other experimental procedures remained the same as the other groups. For monoclonal antibody supplementation experiments (Fig. 7A-D), the indicated mice were injected with 100 μl PBS containing 4 μg of TyTx4 or TyTx11 via retro-orbital injections 30 minutes before the typhoid toxin challenge. TyTx4 and TyTx11 IgGs were generated and characterized previously [15, 17]. Two micrograms of purified recombinant typhoid toxin carrying a histidine hexamer on CdtB were administered via retro-orbital injections when a lethal-dose typhoid toxin challenge is indicated. Changes in the behavior, weight, and survival of the toxin-injected mice were closely monitored as previously described [11, 14-16, 47]. Balance beam tests were performed to evaluate toxin-mediated upper motor function deficits, as previously described [13, 14]. To assess the toxicities towards immune cells, blood samples were collected by submandibular bleeding in Microtainer tubes coated with EDTA as an anticoagulant (BD Biosciences), kept at room temperature, and analyzed within 2 hrs after blood collection using a Hemavet 950FS hematology analyzer (Drew Scientific). To prepare plasma samples, whole blood samples collected into EDTA-treated Microtainer tubes were centrifuged for 10 minutes at 1,000 x g to remove cells.

### Measurement of antibody responses specific to typhoid toxin or PltB

Indicated antibody levels in the plasma of vaccinated mice were examined using a direct enzyme-linked immunosorbent assay (ELISA). 96-well plates (Costar) were coated with 50 ng of purified typhoid toxoid or tagless PltB pentamer in 100 μl plate-coating buffer (50 mM carbonate-bicarbonate buffer, pH 9.6), and incubated overnight at 4 °C. Wells were washed with PBS containing 0.05% Tween 20 and blocked with PBS containing 1% BSA for 1 h at 37 °C. Two-fold serial dilution of the plasma contained in 50 μl PBS/0.05% Tween 20/0.5% BSA was added to each well and incubated for 2 hr at 37 °C. After washing, bound antibodies were detected with horseradish peroxidase (HRP)-conjugated anti-mouse immunoglobulin IgG (Southern Biotech) at a 1:10,000 dilution in PBS/0.05% Tween/0.5% BSA. Wells were then incubated with an HRP substrate, tetramethylbenzidine (Sigma), for 10-30 min, and the reaction was stopped by the addition of 100 μl of 1M H_3_PO_4_. The results were assessed by reading a Tecan Infinite 200 Pro microplate reader. Relative serum antibody levels were determined by standard end-point ELISA titration.

### Immunofluorescence staining

#### Cultured cell staining

Immunofluorescence assays were performed to evaluate the neutralizing function of maternal antibodies in the context of *S*. Typhi infection. In brief, Henle-407 human intestinal epithelial cells were cultured in DMEM high glucose + 10% FBS (HyClone) and kept at 37°C in a cell culture incubator with 5% CO_2_. Cells were seeded on coverslips placed in 24-well culture plates and incubated overnight. On the following day, cells were infected with either wild-type *S*. Typhi carrying the pKan plasmid or *S*. Typhi CdtB^H160Q^ carrying the pKan plasmid at a multiplicity of infection of 30 for 1 hour in HBSS (Invitrogen), treated with 100 μg/mL gentamicin to kill extracellular bacteria for 45 min, and washed with PBS. Infected cells were then incubated in the complete cell culture medium containing 50 μg/mL kanamycin and 10 μg/mL gentamicin for 24 hours in the presence and absence of indicated mouse plasma samples. After incubation for 24 hrs, cells were fixed with 1% paraformaldehyde (PFA) for 10 min, washed with PBS, and permeabilized in PBS containing 50 mM NH_4_Cl, 0.2% Triton X-100, and 3% BSA for 30 min. The slides were stained with indicated antibodies (anti-*S*. Typhi antibody, 1:2000, anti-Flag M2 antibody, Sigma #F1804, 1:2000, anti-phospho-histone H2AX antibody, Thermo Fisher #LF-PA0025, 1:1000) for 2 hrs at room temperature, counterstained with 4′,6-diamidino-2-phenylindole (DAPI), mounted using ProLong antifade mounting solution (Molecular Probes, Thermo Fisher Scientific), and viewed. Immunofluorescence images were acquired with a Leica DMI6000B/DFC340 FX fluorescence microscope system. The fluorescence signal intensity of images was quantified using the measure function of ImageJ (National Institutes of Health, USA) after subtracting the background. Recorded images were merged using the ImageJ merge channels function and processed further with Adobe Photoshop to adjust the brightness and contrast equally for all recordings.

#### Murine tissue staining

Fluorescence microscopy was performed to quantify the FcRn expressions on the yolk sac of F1 mice harvested on gestation day 19 and small intestines of F2 mice on day 7 after birth. The mice were anesthetized with isoflurane, followed by sacrifice– perfusion, which was conducted by sequentially administering 50 mL of each 10% sucrose and 4% paraformaldehyde. The indicated placenta with yolk sac and intestine samples were extracted and fixed in 4% paraformaldehyde for 24 hours at 4 °C. Tissues were washed with PBS and immersed in 30% sucrose solution overnight for cryoprotection. These tissues were trimmed into smaller pieces and placed in cassettes for Tissue-Tek optimum cutting temperature (OCT) embedding. The embedded tissues were flash-frozen in isopentane cooled to –80 °C. Cryosections of frozen tissue samples were cut to be 8 μm thick and stored at −80 °C until staining. Frozen tissue sections were fixed, washed with PBST (PBS with 0.1% tween 20), blocked in 3% BSA/PBS for 30 min, and incubated with goat anti-mouse FcRn (AF6775, R&D Systems; 1:100) or goat anti-human podocalyxin (AF1658, R&D Systems; 1:100) overnight at 4 °C. Then sections were washed with PBST and incubated with fluorochrome-conjugated secondary antibodies (Molecular Probes) for 1 hour at room temperature in the dark. The nuclei were counterstained with DAPI for 10 minutes. The slides were mounted in an antifade mounting solution (Electron Microscopy Sciences) and viewed. The fluorescence signal intensity was quantified as corrected total cell fluorescence (CTCF) [CTCF = integrated density - (area of total cell x mean fluorescence of background)] using ImageJ software. For each group, a minimum of 10-20 images were taken and the results of two independent experiments were plotted as bar graphs. GraphPad Prism software was used to test the statistical significance of the data.

### Nanopore sequencing

For metagenomics, we isolated and sequenced DNAs of the small intestine contents via a nanopore sequencing and analysis pipeline previously established with modifications [44]. Briefly, fecal samples were collected from the small intestines of ND±Abx and MD±Abx mice, and their genomic DNAs were extracted using QIAamp fast DNA stool kit (Qiagen) without processing additional fragmentation. Genomic DNA quality and quantity were monitored using NanoDrop 2000 spectrophotometer (ThermoFisher Scientific) and Qubit 4 fluorometer (ThermoFisher Scientific). Microbiome DNA library was prepared using an SQK-LSK110 ligation sequencing kit [Oxford Nanopore Technologies (ONT)], followed by sequencing with a MinION Flongle flow cell (R9.4.1) using MinKNOW v21.06.0 (ONT). The raw reads were base-called using Guppy v5.0.11 (ONT), followed by filtering the reads with a minimum q score >7. The filtered reads were processed with WIMP workflow via EPI2ME v3.4.1 (ONT) for microbiome analysis.

### Quantification and statistical analysis

Data were tested for statistical significance with the GraphPad Prism software. The number of replicates for each experiment and the statistical test performed are indicated in the figure legends. Image analysis and quantification were performed using ImageJ.

## Data and materials availability

This article/figures includes all data generated during this study. Further information for the data should be directed to and will be fulfilled by the correspondence, Jeongmin Song.

## Acknowledgments

This work was supported in part by NIH AI139625, AI137345, AI141514, and AI141514-03S1 to J.S. The funders had no role in the study design, data collection and analysis, decision to publish, or preparation of the manuscript.

## Author contributions

D. N. conducted experiments shown in Figs 4H-J, 5B-F, 6, and 7E-I. C.A. conducted experiments shown in Figs 1, 2, 3, 4A-G, and 7A-D. Y.-A.Y. made initial observations of maternal antibody-mediated early-life protection. G. L. conducted experiments shown in Fig. 5A. D. N. and C.A. wrote the manuscript draft. J.S.: conceptualization, funding acquisition, supervision, writing – original draft, writing – review & editing.

---

**Supplementary Table 1. Microbiome analysis data, related to Fig. 5A**

**Table 1. Normal and malnourished isocaloric diets used in this study, related to Fig. 2**

**Table 2. SHIRPA of 5-week-old F2 mice fed with normal and malnourished isocaloric diets, related to Fig. 2**.

**Table 3. Fat and protein supplementations used in the study, related to Fig. 7**.

